# Brewery Waste as a Sustainable Protein Source for the Banded Cricket (*Gryllodes sigillatus*)

**DOI:** 10.1101/2024.08.21.609009

**Authors:** S.Y. Kasdorf, M.J. Muzzatti, F. Haider, S.M. Bertram, H.A. MacMillan

## Abstract

Crickets, like other edible insects, can convert organic by-products of the food and agricultural industries into high-value protein. Waste products high in protein like brewer’s spent grain and brewer’s spent yeast are particularly attractive replacements for unsustainable protein sources in cricket feed like fishmeal or soy. Such replacement will only be advantageous, however, if feeding on these waste products does not impact, or only minimally impacts, cricket survival, growth, and body composition. In this study, a farmed cricket species, *Gryllodes sigillatus*, was reared in isolation on experimental diets in which fishmeal was wholly or partially replaced with either brewer’s spent grain or brewer’s spent yeast. Cricket survival, development and macromolecular composition were not different across diets. However, wholly replacing fishmeal with brewer’s spent yeast or brewer’s spent grain reduced cricket adult body mass by approximately 16%. To extend these findings toward a farm environment, a second cohort of crickets were reared communally on diets in which fishmeal, and fishmeal and soy (a secondary protein source), were replaced by brewer’s spent grain. We found that in a communal environment, crickets reared on both diets performed equally as well as the control. Therefore, brewing waste products are promising candidates for use as a primary protein source in the feed of *G. sigillatus*. In addition to contributing towards the goals of a circular bioeconomy through the repurposing of waste, the use of brewing waste in cricket feed may have a positive impact of the cricket farming industry as a cost-effective and sustainable alternative to traditional feed.

**Conflict of Interest:** The authors have an ongoing research agreement with Aspire Food Group and Entomo Farms, who produce crickets as food and feed.

**Funding Statement:** This research was funded by Discovery Grants awarded by the Natural Sciences and Engineering Research Council of Canada (NSERC) to H.A.M. (RGPIN-2018-05322) and S.M.B. (RGPIN-2017-06263). Additional support was provided by an NSERC Alliance Grant (568647-21) and a Deep Space Food Challenge grant (22IUCCAR22) from the Canadian Space Agency awarded to both H.A.M. and S.M.B. Equipment used in this study was purchased with support to H.A.M. from the Canadian Foundation for Innovation (project number 37721).

## Introduction

As insect farming becomes an increasingly popular source of protein worldwide, there is interest in improving the economic and environmental sustainability of insect production. Farmed insects are already less environmentally degrading compared to traditional livestock due to reduced land requirements (Alexander *et al*., 2017), water consumption (Miglietta *et al*., 2015), and greenhouse gas emissions (Huis, 2013; Oonincx *et al*., 2015). However, feed remains a challenge for the industry due to the cost and relative unsustainability of traditional feed ingredients high in protein, such as fishmeal (Tyapkova *et al*., 2016) and soy (Sorjonen *et al*., 2019). Due to the capacity of insects to convert by-products of the food and agricultural industries (Van Huis *et al*., 2021), a solution may be replacement of traditional feed components with nutritionally-valuable organic waste products.

Large-scale organic waste products are often environmentally degrading due to gas production during microbial decomposition (Bharat Helkar and Sahoo, 2016). They are also financially burdensome to producers who must pay for their proper disposal. However, recent efforts have been made to repurpose waste from the food and agricultural industries as a means of increasing global sustainability. This process of returning organic by-products to the production chain has been referred to as the circular bioeconomy (Zeko-Pivač *et al*., 2022), and offers an opportunity to recover valuable resources and reduce production costs.

Therefore, the use of organic by-products in insect feed as a replacement for less sustainable products would be beneficial to the insect farming industry, as well as furthering the goals of a circular bioeconomy.

Beer is the fifth most popular beverage worldwide (Rachwał *et al*., 2020). As a result, modern brewing is a massive global industry that annually produces over 1.87 billion hectolitres of beer (Amienyo and Azapagic, 2016) and 36.4 million tonnes of organic waste (Zeko-Pivač *et al*., 2022). This waste is mainly in the form of brewer’s spent grain (BSG), a solid by-product of the mashing process in which malted grain is combined with water to produce wort, and brewer’s spent yeast (BSY), the yeast left over from the fermentation process (Rachwał *et al*., 2020). BSG and BSY are nutritionally valuable due to their high protein content (Mussatto *et al*., 2006; Thiago *et al*., 2014). Therefore, these products are sometimes recycled as cost-effective feed for pig and poultry livestock (Zeko-Pivač *et al*., 2022). There is interest in extending the use of these products to insect farming (Varelas, 2019).

Black soldier flies (Adebayo *et al*., 2021; Bava *et al*., 2019; Chia *et al*., 2020), crickets (Lundy and Parrella, 2015; Miech *et al*., 2016; Sorjonen *et al*., 2019) and mealworms (Oonincx *et al*., 2015; van Broekhoven *et al*., 2015) can be raised on diets containing by-products of the brewing industry (primarily BSG). Many studies that incorporate brewing waste into insect feed provide the by-product as the sole source of nutrition or in some combination with other organic waste products (Kuo and Fisher, 2022), which often results in lower survival and growth rates compared to a control feed (e.g., chicken feed). By contrast, the use of BSG as an alternative to a traditional protein source (soy) within a base feed has yielded mostly comparable results in both *Acheta domesticus* and *Gryllus bimaculatus* crickets (Sorjonen *et al*., 2019). It is therefore plausible that brewing waste can be used to replace traditional protein sources in existing farm feeds with minimal impacts on product yield or quality, and that wholesale replacement of feed with waste products can yield misleading results. However, before replacement of traditional feed components by brewery waste can be performed at scale, these novel diets must be demonstrated to approach or match rates of insect protein production using existing feeds.

In large-scale insect farming, production success is measured by three primary factors: cost (of feed, temperature regulation etc.), product yield and product quality. Yield is mainly dependent on the number of insects harvested (survival rate), the weight of the insects harvested (body weight) and the time needed to reach harvestable size (development rate). Product quality is dependent on the nutritional profile of the insect’s whole body. As diet can significantly impact survival and growth (Adebayo *et al*., 2021; Bava *et al*., 2019; Oonincx *et al*., 2015; Orinda *et al*., 2017; van Broekhoven *et al*., 2015), as well as nutritional composition of edible insects (Chia *et al*., 2020; Jucker *et al*., 2022; Spranghers *et al*., 2017), these factors are all directly relevant to evaluating the suitability of new diet formulations.

The primary aim of our study was to evaluate the capacity of solid brewing waste to replace fishmeal as the primary protein source in feed used to rear *G. sigillatus*. Many insect farming feeds use fishmeal as their primary protein source (Morales-Ramos *et al*., 2020; Ssepuuya *et al*., 2021), but no studies have tested the suitability of brewing waste as a direct replacement for fishmeal in insect feed. Different species can have varied responses to the same rearing substrate (Oonincx *et al*., 2015; Sorjonen *et al*., 2019), so novel waste diets must be explored for individual species prior to their integration. The edible cricket species *Gryllodes sigillatus* is a popular species for farm production (Zielińska *et al*., 2015), and no studies, to our knowledge, have tested the impacts of including brewery waste in *G. sigillatus* diets.

We performed two experiments: one in which the crickets were reared in isolation, and a second in which the crickets were reared in communal bins to simulate the social environment of farming conditions. Our results indicate that brewing waste contains the necessary protein requirements for *G. sigillatus* and is a suitable replacement for fishmeal (and soy) in cricket feed. Therefore, the use of brewing waste as a protein source to rear edible crickets serves as a prime example of how the constructive utilization of waste products can be integrated into modern agriculture and food production, and the circular bioeconomy.

## Materials and Methods

### Raw Materials

BSG and BSY (containing no hops to avoid a bitter flavour) were kindly donated by Stray Dog Brewing Company in Ottawa, Ontario. Both waste products were dried in a drying oven at 30°C for seven days, then ground using a SmartGrind™ electric coffee and spice grinder (Black & Decker model: CBG100SC) followed by manual grinding using a mortar and pestle. The ground products were passed through a 510 nm steel sieve and stored at -80°C. The fishmeal and all other components of the cricket feed were obtained dried and ground from Campbellford Farm Supply in Campbellford, ON, Canada, (Table 1).

**Table 1.** Analysis output for variables quantified in the isolated rearing experiment.

| Dependent Variable | Model | Factor | Df | F value | p-value |
| --- | --- | --- | --- | --- | --- |
| Survival | Kaplan-Meier Survival Analysis | Protein Source | 4, 108 | NA | 1 |
| Days to Adulthood (Development) | Kaplan-Meier Survival Analysis | Protein Source | 4, 108 | NA | 0.200 |
| Final Adult Mass | Two-Way ANOVA | **Protein Source** | **4, 104** | **2.51** | **0.030** |
|  |  | **Sex** | **2, 104** | **43.38** | **<0.001** |
|  |  | Protein Source * Sex | 6, 104 | 0.50 | 0.128 |
| Final Adult Size | Two-Way ANOVA | **Protein Source** | **4, 104** | **2.72** | **0.028** |
|  |  | **Sex** | **2, 104** | **68.85** | **<0.001** |
|  |  | Protein Source * Sex | 6, 104 | 0.92 | 0.147 |
| Adult Whole-Body Protein Content | Two-Way ANOVA | Protein Source | 4, 101 | 1.74 | 0.159 |
|  |  | **Sex** | **2, 101** | **60.53** | **<0.001** |
|  |  | Protein Source * Sex | 6, 101 | 0.03 | 0.06 |
| Adult Whole-Body Carbohydrate Content | Two-Way ANOVA | Protein Source | 4, 101 | 1.55 | 0.311 |
|  |  | Sex | 2, 101 | 4.21 | 0.079 |
|  |  | Protein Source * Sex | 6, 101 | 1.38 | 0.086 |
| Adult Whole-Body Lipid Content | Two-Way ANOVA | Protein Source | 4, 101 | 0.66 | 0.354 |
|  |  | **Sex** | **2, 101** | **24.52** | **<0.001** |
|  |  | **Protein Source \* Sex** | **6, 101** | **3.44** | **0.008** |

### Isolated Rearing Experiment

To monitor and distinguish between individuals over time, this experiment used individual rearing containers. Five diets were used to evaluate the capacity of both brewery waste products (BSG and BSY) to replace fishmeal as the primary protein source in cricket feed: the control (an existing diet containing fishmeal), and four experimental diets in which the fishmeal was partially or wholly replaced with one of the waste products.

In order to ensure that each diet had similar protein levels, replacement of fishmeal (68% protein, dry weight (d.w.)) was performed by protein content rather than by mass. A Bradford assay (Biorad, Hercules, USA) was used to determine the protein content of BSG (27.6% d.w.) and BSY (44.7% d.w.). Diet creation was based off an existing cricket feed formula containing corn, soy, linseed, fishmeal and a micronutrient supplement. All diet components were mixed using a food processor and stored at -80°C until use.

The isolated rearing experiment was conducted using *Gryllodes sigillatus* nymphs obtained as eggs from Entomo Farms in Norwood, Ontario. As newly emerged nymphs are very delicate, the nymphs were reared communally in rectangular plastic bins (22.9 L x 15.2 W x 5.2 H cm) on a standard farm feed for two-weeks prior to the experiment to avoid deaths unrelated to the experimental treatments. The bins were maintained in an incubator at 32°C, a 14L:10D photoperiod and approximately 50% relative humidity. The two weeks old nymphs were transferred to individual plastic rearing cups (3.5 oz condiment cups) with mesh in the lids for aeration. The cups included a piece of egg carton for shelter and a microcentrifuge tube of water plugged with cotton. Twenty-seven nymphs were randomly assigned to each diet and fed *ad libitum*. All water tubes were refilled thrice weekly when checking individuals for survival, determining their sex, and identifying maturity date; feed was replaced once weekly. All crickets were kept in an incubator at 32°C, a 14L:10D photoperiod and approximately 50% relative humidity for the duration of the four-week experiment. Each cricket’s mass was quantified at the beginning and end of the experiment using an AB135-S scale (Mettler Toledo, Columbus, United States). Body size was quantified by capturing an image of each cricket at the beginning and end of the experiment using a Stemi 508 microscope (ZEISS, Oberkochen, Germany) with an Axiocam 208 colour camera (ZEISS) and ZEN Lite (ZEISS) software. Head width, pronotum width and pronotum length measurements (Figure 1) were obtained from the images using ImageJ v.1.53 software (National Institutes of Health, Bethesda, MD, U.S.A.). Development time was calculated as the number of days between the experiment start date (two weeks post-hatch) and the date the cricket was observed to moult to adulthood.

**Figure 1.**
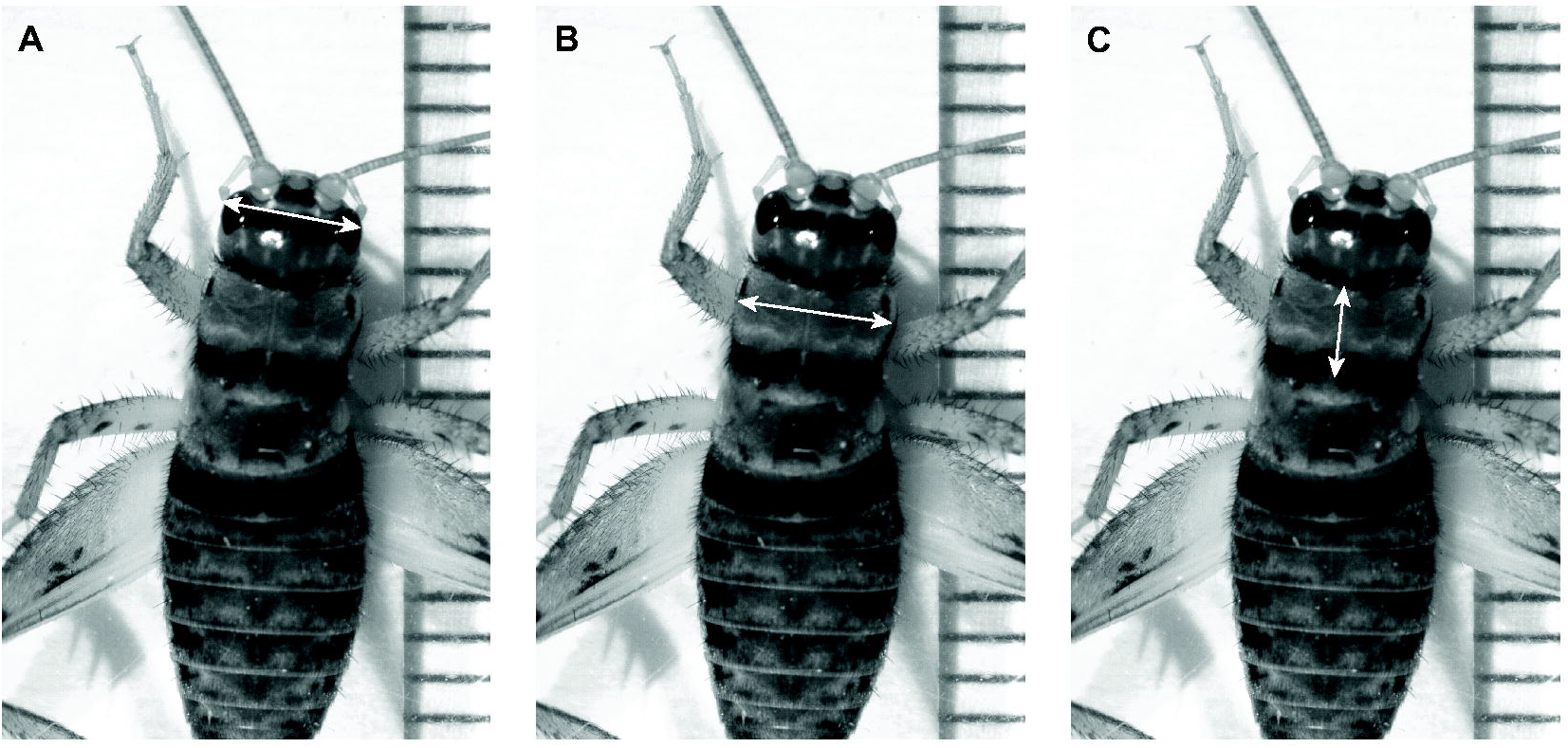
Example measurements for a) head width, b) pronotum width and c) pronotum length.

At the end of the experimental trial, crickets were snap frozen with liquid nitrogen and stored in individual microcentrifuge tubes at -80°C. Tissue energy reserves (protein, carbohydrate, and lipid content) were measured from the whole cricket bodies (n = 20-24 samples per diet). Crickets were homogenized in ice-cold buffer (0.1 M Tris-HCl (pH 8.0), 0.2% Triton X-100, 153 μM MgSO_4_, and 0.1 mM PMSF) in a Fisherbrand™ Bead Mill 24 Homogenizer (Fisher Scientific, Ottawa ON, Canada). For lipid estimation raw homogenate was used and for protein and carbohydrate estimation the homogenate was centrifuged at 3000x g at 4°C and the supernatant was used. Macromolecule estimation was done by optimizing the colorimetric methods from Haider *et al*. (2018). Briefly, lipids were extracted using chloroform-methanol mixture (Folch method) and quantified using phospho-vanillin method. Protein was measured using Bradford assay (Bio-Rad, Hercules, USA). Total carbohydrate was quantified using sulphuric acid-phenol method. All colorimetric samples were analyzed in triplicate.

### Isolated Rearing Experiment Statistical Analyses

All statistical analyses were performed using R (v4.2.3; (R Core Team, 2024)) software. Principal component analyses were used to obtain one value representative of overall body size using the three body size measurements (head width, pronotum width, pronotum length) obtained for the crickets at the beginning and end of the experimental period (Visanuvimol and Bertram, 2011). The PC1 size values accounted for 97% (eigenvalue = 2.91) and 96% (eigenvalue = 2.87) of the variation in cricket size at the beginning and end of the experiment, respectfully. A Kaplan-Meier survival analysis model and log-rank test was used to determine the effect of protein source on survival and development (time to adulthood). Two-way ANOVAs (e.g., final body mass ∼ protein source x sex) were used to determine the effect of protein source on final body mass, final body size and macromolecule content. Tukey’s HSD tests were used to clarify differences among treatment groups for ANOVA models. The individuals that escaped (n = 7) were not included in final body mass, final body size or macromolecule content analyses due to a lack of end-point data.

### Communal Rearing Experiment

Crickets were reared in communal bins to account for social stressors typically experienced during commercial production. In addition to the control (an existing diet containing fishmeal), we tested two experimental diets: fishmeal wholly replaced by spent grain, and fishmeal and soy (a secondary protein source) wholly replaced by spent grain.

Similar to the isolated rearing experiment, replacement of fishmeal (68% protein d.w.) and soy (48% protein d.w., (Yamka *et al*., 2003)) by spent grain (27.6% protein d.w.) was performed by protein content rather than by mass to a maximum total spent grain content of ∼78% on the BSG (soy replaced) diet. Diet creation was based off an existing cricket feed formula containing corn, soy, fishmeal, linseed and a micronutrient supplement. All components were mixed using a food processor and stored at -80°C.

The communal rearing experiment was conducted using an identical rearing protocol to that used for experiment one (section 2.2), with the exception that the two-week-old nymphs were transferred to small communal bins (22.9L x 15.2W x 15.2H cm) instead of individual rearing cups at the beginning of the experimental period (n = 15 individuals per bin, and three bins per diet).

The mass of each cricket was individually measured weekly using an AB135-S scale (Mettler Toledo, Columbus, United States). Survival was quantified by recording the number of living individuals in each bin thrice weekly. Development was monitored by counting the number of crickets in each bin that had moulted to adulthood thrice weekly.

In order to monitor feed consumption, feed was weighed prior to distribution. The leftovers were collected at the end of each week, dried at ∼30°C for at least seven days, then the frass was removed and the contents were reweighed. Feed consumption was defined as the difference between the distributed dry feed and the dried leftovers. The feed conversion ratio (FCR) was calculated with the following equation:

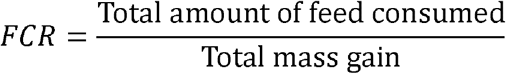

### Communal Rearing Experiment Statistical Analyses

All statistical analyses were performed using R (v4.2.3; (R Core Team, 2024)) software. A Shapiro-Wilks normality test was used to test normality. Mixed effects models (e.g., body mass ∼ protein source x day of experiment + (1|Bin)) were used to evaluate the effect of protein source over time (day of experiment) on survival rate, proportion of adults in a bin (development time) and body mass. A one-way ANOVA was used to evaluate the effect of protein source on feed conversion ratio (FCR).

## Results

### Isolated Rearing Experiment

Dietary protein source impacted adult body mass (F_4,104_ = 2.51, p = 0.030, Figure 2a) and adult body size (F_4, 104_ = 2.72, p=0.028, Figure 2b) of *G. sigillatus* reared in an isolated environment. Specifically, crickets reared on the diet in which fishmeal was wholly replaced with BSY weighed about 18.6% less (184 ± 50 mg) than the control crickets (226 ± 61 mg). Crickets reared on the diet in which fishmeal was wholly replaced with BSG were smaller (PC1 size: -0.355 ± 0.318) than the control crickets (PC1 size: 0.717 ± 0.350). However, crickets reared on the diets in which fishmeal was partially replaced with BSY or BSG did not differ significantly from the control or each other. Survival and development time (Figure 2c) were equal across all protein sources. The whole-body composition (protein, carbohydrate and lipid content) of the crickets was also unaffected by protein source (Figure 3). Sex explained a significant amount of the variation in adult body mass (F_2, 104_ =43.38, p<0.001), adult body size (F_2, 104_ =68.85, p<0.001), whole-body protein content (F_2, 101_ = 60.53, p<0.001) and whole-body lipid content (F_2, 101_ = 24.52, p<0.001). There was no significant interaction between protein source and sex for any of the variables, with the exception of whole-body lipid content (F_6, 101_ = 3.44, p=0.008). Models and statistical measurements are given in Table 1.

**Figure 2.**
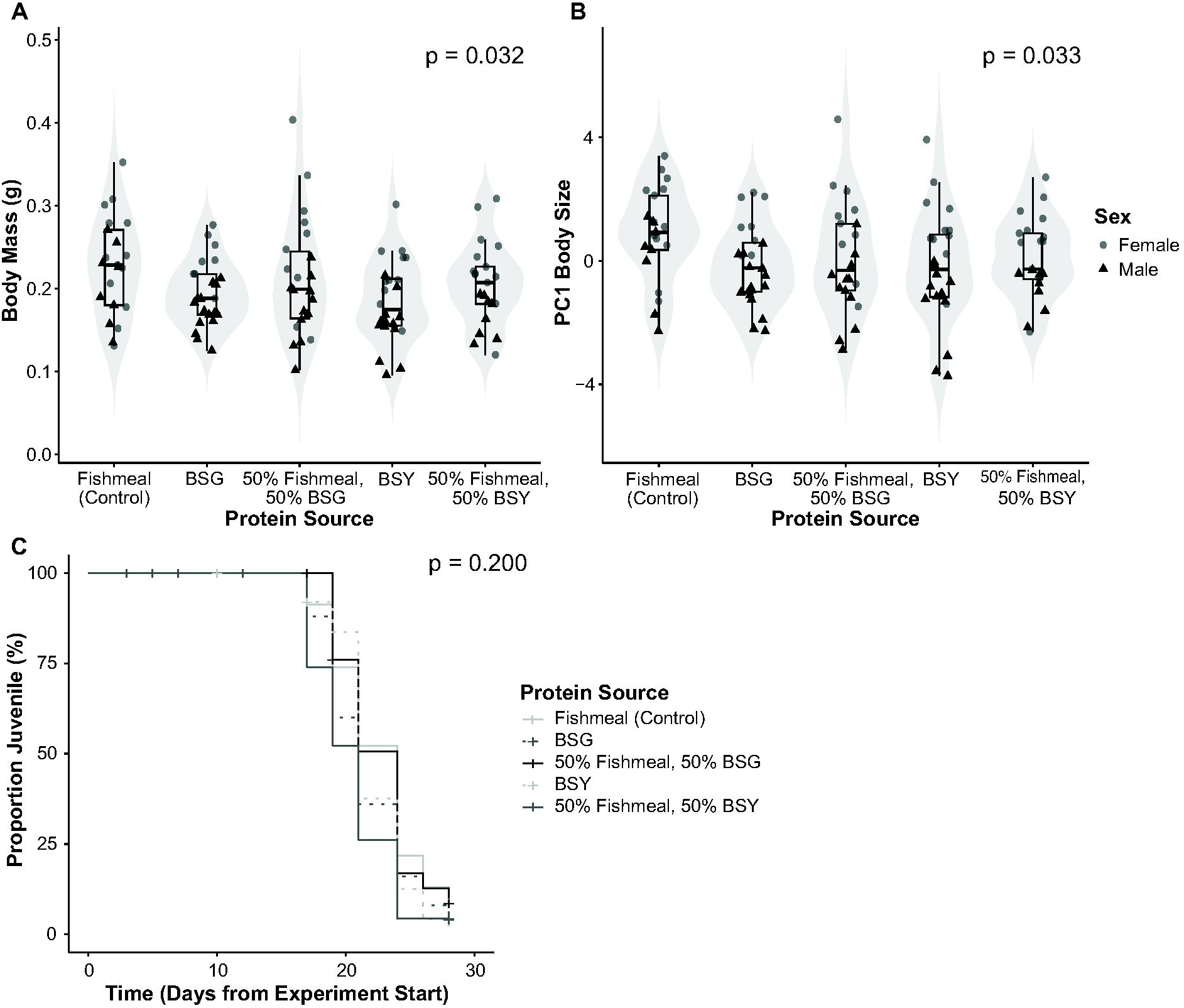
a) Adult cricket mass, b) adult size PC1 and c) proportion of juveniles over time (development time) of crickets reared on each of the experimental diets in the isolated rearing experiment. Each data point represents one individual.

**Figure 3.**
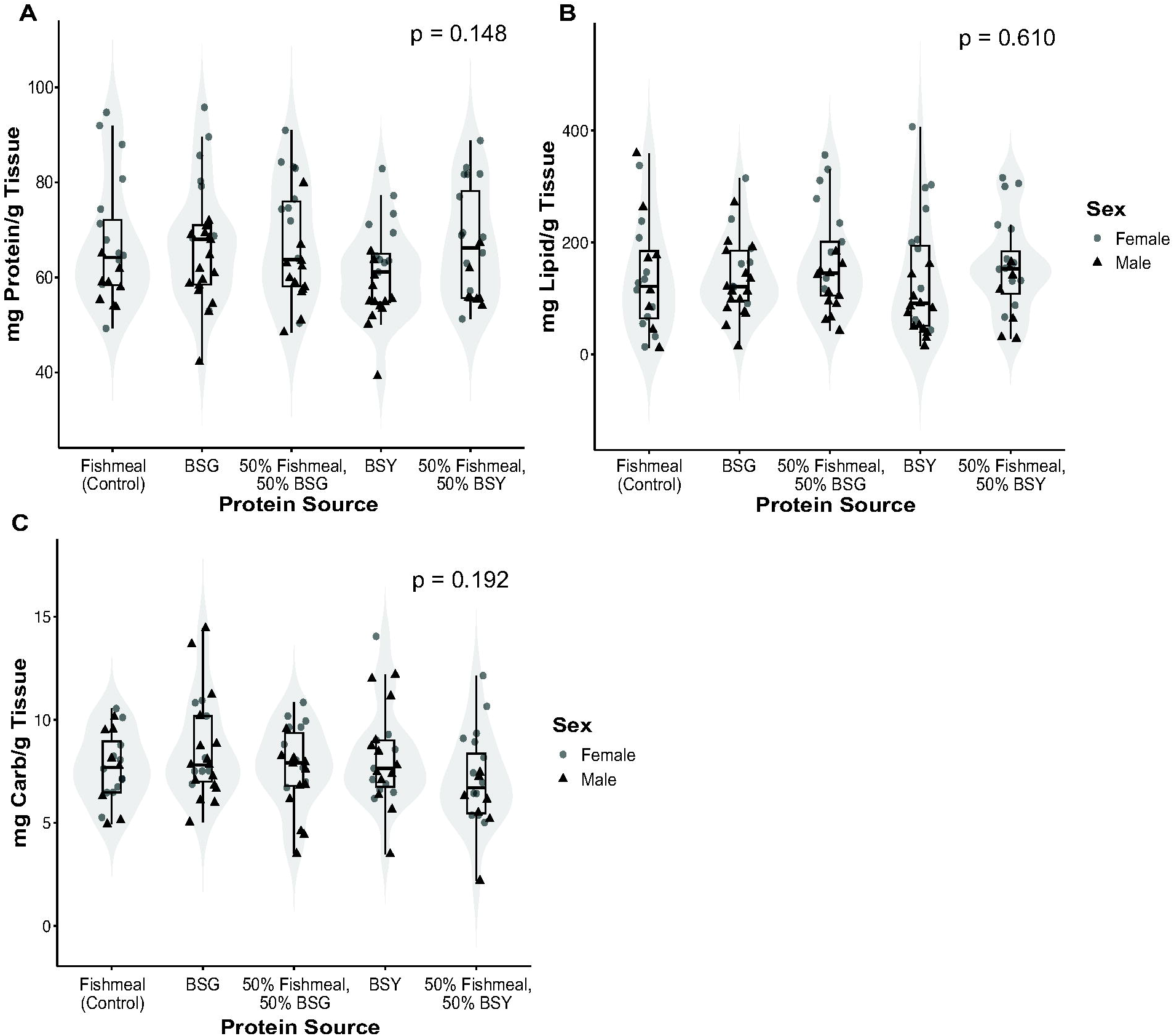
a) Protein, b) lipid and c) carbohydrate content of the cricket whole body reared on each of the experimental diets in the isolated rearing experiment. Each data point represents one individual.

### Communal Rearing Experiment

Crickets reared on all diets performed equally well. Dietary protein source did not have a significant impact on body mass (Figure 4a), proportion individuals that had reached adulthood over time (Figure 4b), survival (Figure 4c) or feed conversion ratio (FCR; Figure 4d) for crickets reared in communal environments. Models and statistical measurements are given in Table 2.

**Table 2.** Analysis output for variables quantified in the communal rearing experiment.

| Dependent Variable | Model | Factor | F value | p-value |
| --- | --- | --- | --- | --- |
| **Survival Rate (over time)** | Mixed Effects Model | Protein Source (fixed) | 0.55 | 0.604 |
|  |  | **Day of Experiment (fixed)** | **8.25** | **<0.001** |
|  |  | Protein Source * Day of Experiment | 0.53 | 0.960 |
| **Proportion of Juveniles (over time)** | Mixed Effects Model | Protein Source (fixed) | 0.01 | 0.334 |
|  |  | **Day of Experiment (fixed)** | **561.14** | **<0.001** |
|  |  | Protein Source * Day of Experiment | 0.26 | 0.247 |
| **Body Mass (over time)** | Mixed Effects Model | Protein Source (fixed) | 1.07 | 0.392 |
|  |  | **Day of Experiment (fixed)** | **5234.17** | **<0.001** |
|  |  | Protein Source * Day of Experiment | 0.344 | 0.709 |
| **Feed Conversion Ratio (FCR)** | One-Way ANOVA | Protein Source | 3.546 | 0.096 |

**Figure 4.**
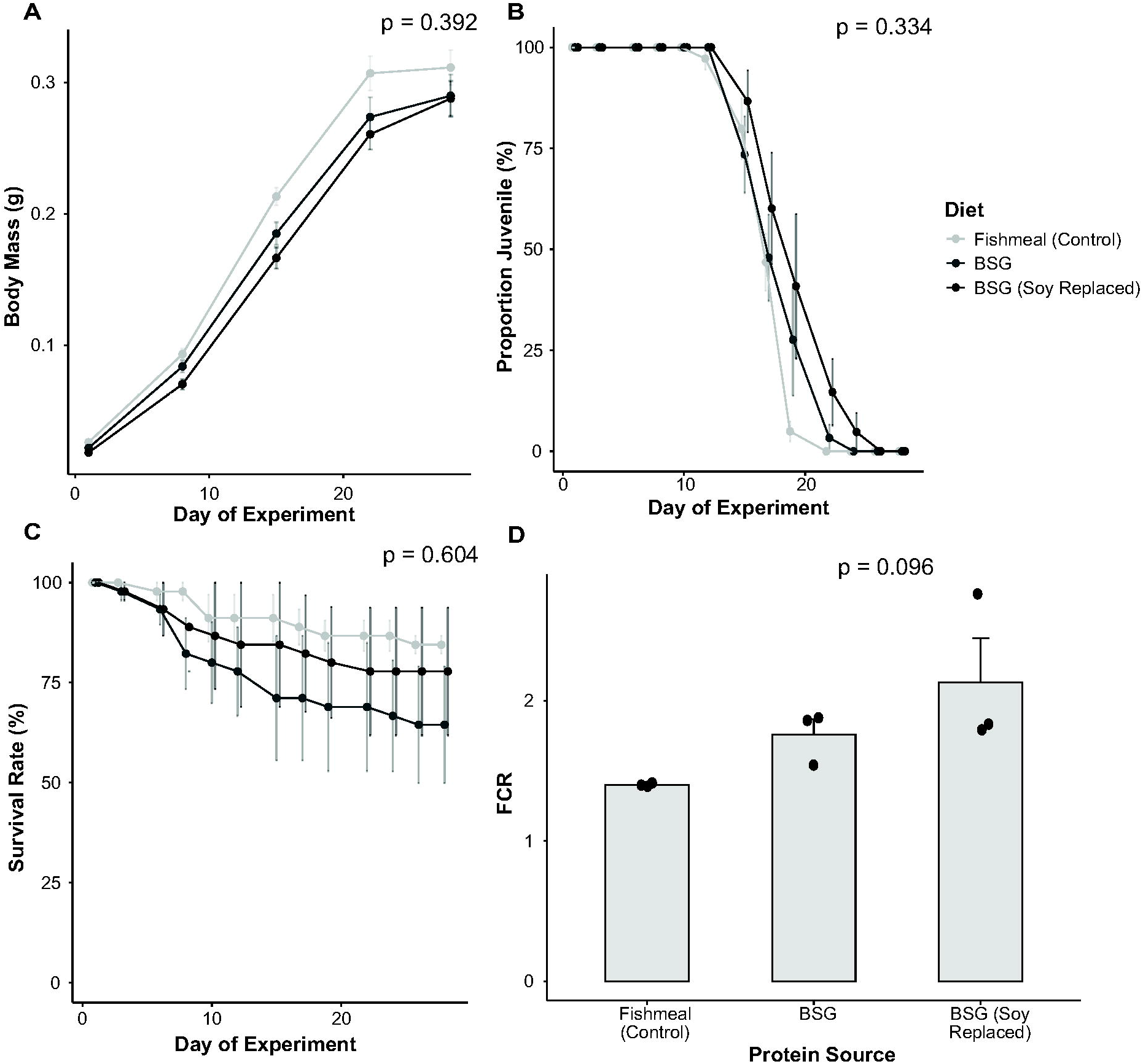
a) Average body mass, b) average proportion of juveniles (development), c) average survival rate, and d) feed conversion ratio (FRC) by protein source for crickets reared in the communal rearing experiment. Error bars represent standard error.

## Discussion

Our study is the first to investigate the use of solid brewing waste (BSG or BSY) as a whole or partial replacement for fishmeal in cricket feed. To our knowledge, it is also the first to investigate the use of these products for the edible cricket species *Gryllodes sigillatus*, which is an established species for production (Zielińska *et al*., 2015) that rarely features in the insects as food and feed literature. We found that both BSG and BSY are viable options for further exploration as replacements for fishmeal in the feed of *G. sigillatus*. These findings have the potential to create a positive impact on the cricket farming industry by providing access to a more sustainable and cost-effective source of feed, as well as contributing towards the constructive utilization of waste products for global sustainability.

In isolated rearing conditions, replacing fishmeal with either BSG or BSY did not impact the whole-body protein (Figure 3a), lipid (Figure 3b) or carbohydrate content (Figure 3c) of the crickets; the macromolecular nutritional quality was equal to that of control crickets reared on a standard farm feed containing fishmeal. Survival averaged ∼91% for all treatments; our higher than average rates of survival (15-50%; Dobermann *et al*., 2019; Jucker *et al*., 2022; Miech *et al*., 2016; Oonincx *et al*., 2015) may be attributed to the isolated rearing conditions, which eliminates competition among individuals in dense social environments (Gutiérrez *et al*., 2020) Also, the nymphs were placed on the treatment diets after two-weeks rather than at hatching, which likely promoted higher survival due to early-life access to high-quality nutrition (Dobermann *et al*., 2019).

The diet in which fishmeal was fully replaced with brewing waste resulted in a 13.7% reduction in adult body size and a 18.6% reduction in mass for crickets in isolation compared to the control crickets. This mass difference was only significant for the BSY diet. However, while small trends of reduced mass may be nonsignificant at the experimental level, they have the potential to be highly negatively impactful at a commercial scale, as cricket mass is a large contributor to farm yields. The significant reduction in cricket adult body mass observed on the BSY diet may be a result of a lack of necessary amino acids. Fishmeal is a rich source of all ten essential amino acids (EAAs) (Cho and Kim, 2011), and attempts to replace fishmeal in aquaculture with cheaper (often plant-based) protein sources often lead to deficiencies in one or more EAAs, particularly methionine and lysine (Nunes *et al*., 2014).

Indeed, BSY contains low levels of both methionine and arginine (Podpora *et al*., 2016), relative to fishmeal. Dietary EAAs are critical for building tissues (Cohen, 2015), and so limited availability could negatively impact cricket growth. Body weight could potentially be recovered by supplementing a BSY diet with processed amino acids. Supplementation of EAAs to mass reared insects is just beginning to garner attention. While supplementing plant-based diets with crystalline amino acids improved yellow catfish (Wang *et al*., 2022) and broiler chickens (Hilliar *et al*., 2020) yields, crystalline lysine supplements did not influence growth or development of black soldier fly larvae (Koethe *et al*., 2022).

Given the success of the individual-level experiment where crickets were reared in isolation, we conducted a follow-up experiment using communal rearing bins. Communal rearing accounts for the effect of social interactions potentially experienced in a farm environment. This communal experiment included the full-replacement BSG diet due to its success in the previous individual experiment, and excluded BSY because BSG makes up 85% of the waste produced by the brewing industry (Rachwał *et al*., 2020). Further, we aimed to determine how much BSG could be added to the diet without any negatively impacting cricket production. We therefore also included a diet in which both protein and soy (a secondary protein source) were replaced with BSG (77.72% BSG by mass).

In the communal experiment, the crickets reared on the diet in which fishmeal was fully replaced with BSG, and the diet in which both fishmeal and soy were replaced with BSG, had a 20% and 10% lower survival rate, respectively, compared to crickets reared on the control diet. While this difference was not significant at the scale tested, survival was reduced compared to the isolated rearing experiment. Reduced survival in communal conditions may result from increased cannibalism on the waste diets, which can occur when crickets experience suboptimal nutrition (Gutiérrez *et al*., 2020; Randall *et al*., 2023). Cricket weight also tended to be reduced, and crickets reared on the BSG and BSG (soy replaced) diets were approximately 6.7% and 7.4% lighter respectively, than the control crickets at the close of the experiment. These body mass differences are notably less than the body mass difference we observed in the isolated rearing experiment (13.7%), supporting the idea that cannibalism may be partially making up for the shortcomings of the BSG diet. Previous studies rearing edible insects on diets containing BSG also report decreased growth and survival (Jucker *et al*., 2022; Lundy and Parrella, 2015; Miech *et al*., 2016; Oonincx *et al*., 2015). While direct comparison is difficult due to differences in diet composition and waste inclusion levels, our results are similar to those reported by Sorjonen *et al*., 2019 for *A. domesticus* and *G. bimaculatus* reared communally on a diet in which soy (a primary protein) was replaced with BSG. In Sorjonen *et al*., 2019 study, the *A. domesticus* crickets on the BSG diets had comparable survival but were slightly smaller than those reared on the control. There were also differences in the feed conversion efficiency (FCR) of crickets reared on the different diets. The crickets reared on the control diet had an FCR of approximately 1.40. Similar values in the range of 1.34-1.50 have been previously reported for *A. domesticus* (Bawa *et al*., 2020; Lundy and Parrella, 2015; Vaga *et al*., 2020). The FCR increased to 1.76 and 2.13 for the BSG and BSG (soy replaced) diets, respectively, which indicates that the nutrient assimilation efficiency decreases with the addition of brewery waste.

Fiber content could also explain the reduced adult weight and increased FCR on the BSG diets. BSG is high in undigestible polysaccharides such as cellulose, hemicellulose and lignin, which together comprise up to 50% of the product dry weight (Aliyu and Bala, 2011; Ikram *et al*., 2017). High fiber may also limit the availability of other dietary components such as protein and digestible carbohydrates, which could negatively impact growth. For example, *A. domesticus* suffers from reduced body mass when reared on waste diets with higher fiber content (Lundy and Parrella, 2015). This effect was mainly attributed to an inability to access adequate protein and other digestible materials, but protein content was also naturally reduced on high fibre diets, making a cause-and-effect understanding elusive. If fibre content interferes with protein accessibility, the nutritional value of spent grain could potentially be increased using a form of solid state fermentation, a means of degrading lignocellulosic materials (Eliopoulos *et al*., 2022).

The trend toward a decrease in survival of crickets raised communally on BSG diets relative to the control may reflect the influences of a social environment. Interactions with other individuals can result in both cannibalism and competition, which impact survival and growth, particularly on sub-optimal or challenging diets (Gutiérrez *et al*., 2020). Notably, there was also considerable variation in survival among the bins of crickets on the same experimental diet in the communal rearing experiment; this variation was lower in the control group, and the variation was also small for crickets reared individually on the same experimental diet (isolated rearing experiment). Therefore, increased variation in the brewery waste treatments (and its impacts on performance) might result from the interactive effects of the novel diet and the dense social environment, which can influence both the behaviour and life history traits of an insect (Gutiérrez *et al*., 2020; Lihoreau *et al*., 2015). In addition, insects reared on the brewery waste treatments have yet to adjust to the novel diets, and performance on novel diets can increase over successive generations (Ekesi *et al*., 2007; Santos-Garcia *et al*., 2020). This improvement over multiple generations has been tentatively attributed to factors such as modification in the composition of the gut microbiome to better assimilate nutrients (Santos-Garcia *et al*., 2020). As a result, the stress of being introduced to novel diets may have been more acutely experienced in the communal rearing environment. A logical next step in the evaluation of brewer’s waste as a replacement for fishmeal would be analyzing performance over successive generations to evaluate whether among group variation decreases and performance increases as crickets become “accustomed” to the brewery waste diet. A multiple generational study would also enable us to ascertain other factors, like reproductive success, that are important prior to integration of brewery waste as a protein source. Diet can influence factors like fecundity and mating behaviour in insects (Geister *et al*., 2008; Harrison *et al*., 2014; Magara *et al*., 2019). Furthermore, the performance of offspring can be impacted by the diet of the parents due to maternal effects (Bernardo, 1996; Kyneb and Toft, 2006). Therefore, the impact of brewery waste diets on reproductive metrics (e.g., fecundity, egg hatchability, offspring viability) and offspring viability is crucial.

Despite the decreases in yield (survival and growth) of crickets reared on diets containing brewery waste observed in our study, the integration of BSG or BSY into commercial cricket feed could still be economically beneficial for farms. In 2022, the cost of fishmeal from Peru (the largest producer and exporter of fishmeal) was $1,642 USD per metric ton, an increase of 10% from the previous year (USDA, 2023). Prices can only be expected to continue to rise with growing demand (Macusi *et al*., 2023). In contrast, spent grain and brewer’s residual yeast are waste products that could potentially be obtained by farms at low, or potentially no cost from local breweries. Therefore, brewing waste, particularly BSG, offers a promising alternative to traditional protein sources for future exploration in cricket feed.

## Acknowledgements

Our thanks to the individuals at Entomo Farms and Stray Dog Brewing Company for their continued support and contributions to this work.

## Author Contributions

Experiment conceptualization and design were performed by S.Y.K., S.M.B, and H.A.M. Rearing experiments were performed by S.Y.K. with help from M.J.M., and macromolecular analysis of cricket whole-body samples was performed by F.H. Supervision was performed by M.J.M., S.M.B. and H.A.M. Project administration, funding acquisition and resource provision were performed by S.M.B and H.A.M. Data visualization and analysis was performed by S.Y.K. Original draft was written by S.Y.K. All authors contributed to editing and review.

## Supplementary Material

Raw data files and metadata included in submission.

**Table S1**. Isolated rearing cricket performance data.

**Table S2**. Isolated rearing cricket macromolecular composition data. **Table S3**. Communal rearing cricket survival and development data. **Table S4**. Communal rearing cricket body mass data.

**Table S5**. Communal rearing cricket feed consumption data.

